# SE(3)-Equivariant Energy-based Models for End-to-End Protein Folding

**DOI:** 10.1101/2021.06.06.447297

**Authors:** Jiaxiang Wu, Tao Shen, Haidong Lan, Yatao Bian, Junzhou Huang

## Abstract

Accurate prediction of protein structures is critical for understanding the biological function of proteins. Nevertheless, most structure optimization methods are built upon pre-defined statistical energy functions, which may be sub-optimal in formulating the conformation space. In this paper, we propose an end-to-end approach for protein structure optimization, powered by SE(3)-equivariant energy-based models. The conformation space is characterized by a SE(3)-equivariant graph neural network, with substantial modifications to embed the protein-specific domain knowledge. Furthermore, we introduce continuously-annealed Langevin dynamics as a novel sampling algorithm, and demonstrate that such process converges to native protein structures with theoretical guarantees. Extensive experiments indicate that SE(3)-Fold achieves comparable structure optimization accuracy, compared against state-of-the-art baselines, with over 1-2 orders of magnitude speed-up.

## 1 Introduction

During the past few decades, the problem of predicting protein structures from amino-acid sequences has attracted increasing attention. There has been a remarkable progress towards more accurate protein structure prediction, largely owing to more precise inter-residue contact/distance and orientation predictions by deep networks [1, 2, 46, 48, 29, 30, 17]. On the other hand, most structure prediction methods rely on traditional optimization toolkits, including Rosetta [33], I-TASSER [26], and CNS [5]. In CASP14^1^, almost all the top-ranked teams adopted either Rosetta or I-TASSER for protein structure optimization, with the only exception of AlphaFold2 [18]. Such toolkits often involve complicated optimization techniques and multi-stage refinement, which imposes extra difficulties in building an end-to-end system for protein structure optimization. Additionally, most statistics energy functions [23, 24] used in these toolkits are built with human expertise, which may be sub-optimal in formulating the protein conformation space.

There have been a few attempts for fully-differentiable protein structure optimization. In [3], LSTM models are trained to estimate backbone torsion angles, which are then used to recover the geometry of protein backbones. NEMO [16] represents protein structures as internal coordinates and performs iterative refinement by a unrolled Langevin dynamics simulator, guided by a neural energy function. AlphaFold2 [18] employs an attention-based fully-differentiable framework for end-to-end learning from homologous sequences to protein structures, and achieves atomic resolution for the first time. However, critical technical details in AlphaFold2 are still unrevealed at the moment.

On the other hand, several works adopt energy-based models (EBMs) and/or normalizing flow to characterize 3D molecular structures, where the SE(3)-equivariance is (approximately) preserved. The SE(3)-equivariance requirement is critical to guarantee consistent and predictable predictions *w.r.t*. arbitrary 3D rotations and translations. Xu *et al*. [47] combine flow-based and energy-based models to predict the inter-atom distance and then recover 3D structures for small molecules. Shi *et al*. [31] firstly estimate gradients over inter-atom distance, which are invariant to rotations and translations, and then propagate these gradients to 3D coordinates with SE(3)-equivariance preserved.[27] adopts E(n)-equivariant graph neural networks (EGNNs) [28] to model the first-order derivative in a continuous-time flow for molecule generation in 3D. In [8], random rotations are used as data augmentation to encourage (but not guarantee) the rotational invariance of the model for protein rotamer recovery, *e.g*. determining the best side-chain conformation with backbone structures given.

However, it remains unclear how to extend above approaches for efficient protein structure optimization. [47, 31, 27] only focus on small molecules consist of tens of atoms, while proteins are macro-molecules and often involve over thousands of atoms. NEMO [16] does not utilize the coevolution information due to its sequence-based formulation, which is critical for determining protein structures. Additionally, its training process is computationally expensive (about 2 months) and could be sometimes unstable due to back-propagating through long simulations.

In this paper, we propose a SE(3)-equivariant energy-based model for fully-differentiable protein structure optimization, namely SE(3)-Fold. Protein structures are formulated as graphs with amino-acid residues as nodes and inter-residue interactions as edges. We adopt EGNN [28] to iteratively update node embeddings and estimated gradients over 3D coordinates in a SE(3)-equivariant manner. This model is trained to approximate the underlying distribution of protein structures. However, simply applying EGNN models for protein folding is insufficient in capturing various types of inter-residue interactions, and may encounter the stability issue during the sampling process. Hence, we introduce multi-head attentions to better formulate protein structures, and propose the continuously-annealed Langevin dynamics (CALD) algorithm for more stable sampling. Furthermore, we present the theoretical analysis on its convergence, and empirically show that CALD consistently stabilizes the protein structure optimization process.

We evaluate our SE(3)-Fold approach for protein structure optimization on the CATH database [32]. In general, SE(3)-Fold achieves comparable structure prediction accuracy (lDDT-Ca: 0.7928) with the state-of-the-art baseline, trRosetta [48]. By cooperating the multi-head attention mechanism and CALD sampling, we observe consistent improvement over the naive combination of EBM and EGNN models. SE(3)-Fold is GPU-friendly and computationally efficient for both training and sampling, which can speed up the structure optimization process by 1-2 orders of magnitude than trRosetta.

We summarize our major contributions in three folds:

- We propose a fully-differentiable approach for protein structure optimization, which paves the way to end-to-end learning from homologous sequences to protein structures.
- We scale up energy-based models for small molecules to proteins with thousands of atoms via residue-based graph representation, which is efficient in both training and sampling.
- To alleviate the stability issue in sampling-based structure optimization, a novel continuously-annealed Langevin dynamics sampling algorithm is proposed with theoretical guarantee.

## 2 Related Work

### Protein structure optimization

Most methods predict protein structures by minimizing statistics energy functions [23, 24] and/or performing comparative modeling from structural templates [37]. In [2, 48, 30, 17], deep networks are introduced to predict inter-residue contacts, distance, and/or orientations, which are then transformed into additional restraints or differentiable energy terms for structure optimization. Still, traditional structure optimization toolkits, *e.g*. Rosetta [33], are heavily used in these methods for energy minimization. RGN [3] and NEMO [16] directly predict 3D structures with fully-differentiable networks; however, neither of them utilizes the co-evolution information, which leads to inferior quality of predicted structures. AlphaFold2 [18] is the first end-to-end approach to directly build protein structures from co-evolution information embedded in homologous sequences, without the need for Rosetta-like optimization toolkits.

### Energy-based models

These models formulate the probability distribution by parameterizing its unnormalized negative log probability (*i.e*. energy function) or its gradients (*i.e*. score function). Such formulation avoids the restriction on the tractability of normalization constant, allowing various models to be used to depict the probability distribution. Energy-based models have been applied in various domains, including image generation [43, 9, 34, 35, 36], 3D shape pattern generation [44, 45], and molecular conformation prediction [16, 8].

### SE(3)-equivariant models

Due to rotational and translational symmetries in the 3-dimensional space, SE(3)-equivariance is critical for models directly operates on the structural data, *e.g*. 3D point cloud [38] and molecular structures [11, 15]. In [38, 11, 12], spherical harmonics and Clebsch–Gordan coefficients are used to construct SE(3)-equivariant convolutional kernels. Regular representations are used in [7, 10, 15] as an alternative approach for achieving the SE(3)-equivariance. [28] modifies the standard message passing process [13] with an E(n)-equivariant update rule for 3D coordinates.

## 3 Preliminaries

In this section, we briefly review basic concepts of energy-based models and E(n)-equivariant graph neural networks, which are fundamental to our proposed SE(3)-Fold approach.

### 3.1 Energy-based Model

Assuming the dataset 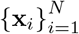 is uniformly sampled from some unknown distribution *p* (**x**), energy-based models (EBMs) aim at approximating its energy function *E* (**x**) = log *p* (**x**) + *C* (where *C* is an arbitrary constant) or score function *s* (**x**) = ∇_x_ log *p* (**x**) with neural networks. Particularly, [34, 35] propose to approximate the score function with denoising score matching (DSM). Each sample **x** is randomly perturbed with an additive Gaussian noise 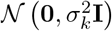, and the perturbed sample is denoted as 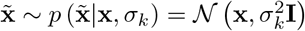. In DSM, *K* random noise levels are used, *i.e.σ*_1_ > *σ*_2_ > *σ*_*K*_, to simultaneously approximate the original distribution and mix among different data modes. The score network, which takes perturbed samples as inputs, is expected to estimate the perturbed data distribution’s log probability’s gradients over perturbed samples. Formally, the optimization objective is given by:

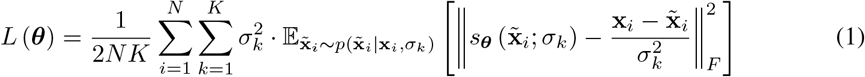

which is the weighted sum of estimation error across all the random noise levels. The above optimization objective aims at minimizing the difference between the data distribution *p* (**x**) and its parameterized counterpart, defined by the score network *s*_***θ***_ (). To sample novel samples from the score network’s corresponding data distribution, one may adopt annealed Langevin dynamics:

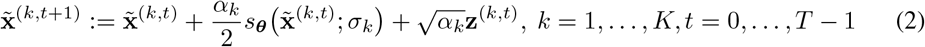

where *α*_*k*_ is the step size for the *k*-th random noise level, and **z**^(*k,t*)^ ∼ 𝒩 (**0, I**). The sampling process starts with *k* = 1, corresponding to the largest standard deviation in the random noise, and then gradually reduces it. Each level’s sampling process is initialized as 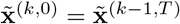, except for the first level, which is sampled from the prior distribution, *i.e*. 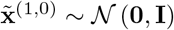.

### 3.2 E(n)-Equivariant Graph Neural Network

#### Equivariance

Let function *f* (·) be a mapping between 𝒳 and 𝒴, and **x** ∈ 𝒳 Consider the abstract group *G* and *g* ∈ *G*, we denote its corresponding transformations as *T*_*g*_ : 𝒳 → 𝒳 and *S*_*g*_ : 𝒴 → 𝒴 The function *f* (·) is considered to be equivariant to *G*, if for any *g* and transformation *T*_*g*_, there exists at least one transformation *S*_*g*_ satisfying:

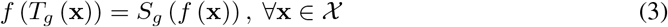

We suggest readers to [11] for a more comprehensive explanation of equivariance.

#### E(n)-Equivariant Graph Neural Network (EGNN)

The EGNN model [28] is designed to guarantee the E(n)-equivariance for point cloud data, where each point is associated with a feature vector **h**_*i*_ ∈ ℝ^*D*^ and a coordinate vector **x**_*i*_ ∈ ℝ^*N*^. The E(n)-equivariance is defined on the *N* -dimensional space, consists of translation, rotation (reflection), and permutation equivariance. The E(n)-equivariant graph convolutional layer (EGCL) modifies the message passing process to ensure the E(n)-equivariance:

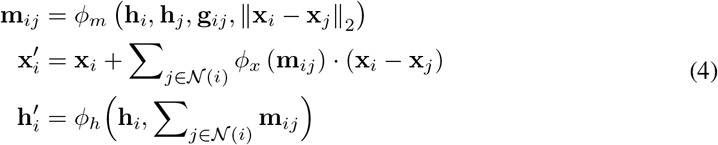

where 𝒩_*i*_ is the *i*-th node’s neighborhood, **g**_*ij*_ is edge features for (*i, j*), and **m**_*ij*_ is the intermediate edge-wise message. Non-linear mappings (*ϕ*_*m*_, *ϕ*_*x*_, and *ϕ*_*h*_) are instantiated as multi-layer perceptron models with Swish activations [25]. Since coordinate vectors are updated by the weighted sum of relative coordinates, and weighting coefficients are E(n)-invariant, the above message passing process is guaranteed to be E(n)-equivariant.

## 4 SE(3)-Fold

In this section, we firstly formulate the protein folding problem from an EBM perspective. Afterwards, we describe how to train the underlying energy-based model, and present a novel sampling algorithm for structure optimization with theoretical guarantee.

### 4.1 Protein Folding: An EBM Perspective

Given an amino-acid sequence of length *L*, the protein folding task aims at determining 3D coordinates of all the atoms for each amino-acid residue. As a simplified form, one may alternatively predict 3D coordinates for backbone atoms (N, C_*α*_, C′, and O) or C_*α*_ atoms only. This is almost sufficient for protein folding, as there are several off-the-shelf tools [14, 20, 40] for determining side-chain conformations from backbone or C_*α*_-trace structures. Hence, in this paper, we only focus on solving C_*α*_-trace or backbone structures, although the proposed method can be easily scaled up for formulating full-atom structures.

For each residue, we take *U* atoms to represent its spatial location, where *U* = 1 for C_*α*_-trace and *U* = 4 for backbone structures^2^. We denote all the considered atoms’ 3D coordinates as **x** ∈ ℝ^3*LU*^, which is the concatenation of each atom’s 3D coordinate **x**_*i,u*_. The amino-acid sequence is denoted as *A*. From the probabilistic perspective, the protein folding task can be viewed as finding the most likely 3D coordinates configuration for the given amino-acid sequence, *i.e*. **x**^∗^ = arg max *p* (**x** | *A*). As the underlying distribution *p* (**x** | *A*) is unknown, we propose to approximate it with an energy-based model, parameterized by a score network *s*_***θ***_ (**x**, *A*). Formally, the score network should satisfy:

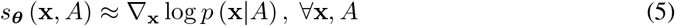

which approximates the log probability’s gradients over 3D coordinates. Once the score network is trained, one may start from randomly initialized 3D coordinates, and gradually refine them via Langevin dynamics. The resulting 3D coordinates are expected to be close to the native structure, if the score network’s corresponding probability distribution is similar with the native one.

### 4.2 SE(3)-Fold: Training

The protein structure can be represented as a graph *G* = (𝒱, ℰ), where vertexes 𝒱 are amino-acid residues, and edges ℰ correspond to inter-residue interactions. This allows us to naturally capture long-range dependencies embedded in protein structures from the graph perspective, which is somewhat difficult for sequence-based formulations. Additionally, we associate each node with per-residue features **h**_*i*_ (*e.g*. amino-acid type and positional encoding), and each edge with pairwise features **g**_*ij*_ (*e.g*. inter-residue distance and orientation predictions) to augment the graph representation. The estimation of log probability’s gradients over 3D coordinates is thus converted into a node regression task, where the SE(3)-equivariance must be preserved. In other words, if the input 3D coordinates are rotated and/or translated by some transformation, then the score network’s outputs must follow the same transformation.

For the score network, we adopt a modified E(n)-equivariant graph neural network as the backbone architecture. Multi-head attention is introduced to capture various types of inter-residue interactions. As each node is now associated with multiple atoms’ 3D coordinates, we update each atom’s 3D coordinates with independent sub-networks to preserve the SE(3)-equivariance. During each message passing process, edge-wise messages are firstly computed to determine attention coefficients:

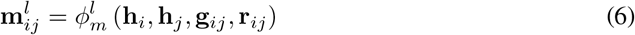

where **r**_*ij*_ encodes the distance between C_*α*_ atoms in the *i*-th and *j*-th residues. The index of attention head is denoted as *l* = 1, …, *L*. Afterwards, query and key embeddings are given by:

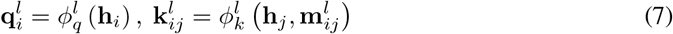

and then passed through a softmax function to compute multi-head attention coefficients:

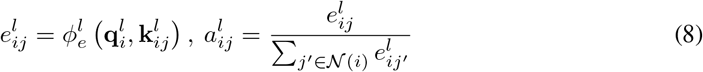

where *𝒩*_*i*_ denotes the *i*-th node’s neighborhood, and 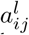 is the attention coefficient for the *l*-th head. Finally, node features and estimated gradients are updated via short-cut connections when possible:

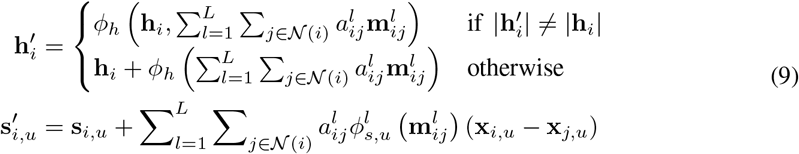

where **s**_*i,u*_ are initialized as all-zeros, and then updated in a residual manner. This is slightly different from [28], which updates 3D coordinates within each EGCL unit, while we keep input 3D structures constant throughout the score network and only update estimated gradients. The detailed implementation of all the above sub-networks can be found in Appendix B.

So far, we have presented the complete workflow of multi-head attention based E(n)-equivariant graph convolutional layer, referred to as MhaEGCL. Our score network consists of multiple stacked MhaEGCL units to increase the model capacity. Similar with denoising score matching [39, 34], the score network is trained to estimate ground-truth gradients over 3D coordinates for the randomly perturbed data distribution 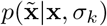. The score network’s estimated gradients are denoted as 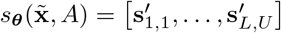 from the last MhaEGCL unit. The loss function is given by:

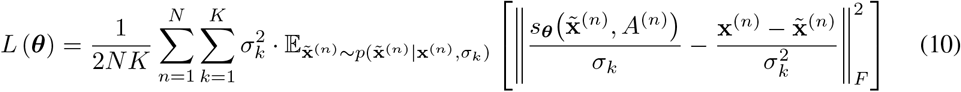

where *A*^(*n*)^ is the *n*-th training sample’s amino-acid sequence. The score network’s outputs are down-scaled by 1*/σ*_*k*_ to match the magnitude of ground-truth gradients at each random noise level, as suggested in [35]. The multiplier 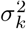 is introduced to approximately keep each random noise level’s equal contribution to the loss function. During each training iteration, native protein structures are perturbed with Gaussian noise at various levels, and then fed into the score network to estimate corresponding gradients over perturbed 3D coordinates.

### 4.3 SE(3)-Fold: Sampling

The score network defines a probability distribution of protein structures *p*_***θ***_ (**x** | *A*) that approximates the native one *p* (**x** | *A*), if the objective function in Eq. (10) is minimized. Therefore, to predict the protein structure from its amino-acid sequence, we start with randomly initialized 3D coordinates, and then apply annealed Langevin dynamics (ALD) [34] for iterative refinement:

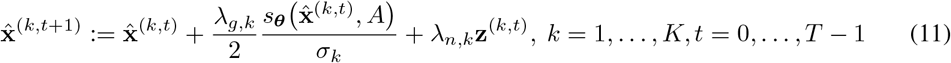

where 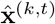 denotes refined 3D coordinates at the *k*-th random noise level’s *t*-th iteration. The multiplier for Gaussian noise is denoted as *λ*_*n,k*_ = *α*_*n*_*σ*_*k*_*/σ*_*K*_, which is proportional to the *k*-th random noise level’s standard deviation. The multiplier for estimated gradients is denoted as 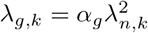, where *α*_*g*_ is newly introduced to control the signal-to-noise ratio.

However, in ALD, the transition between different random noise levels is discontinuous, which may be sub-optimal to the convergence behavior. Therefore, we propose a continuously-annealed Langevin dynamics (CALD) sampling process to smoothly transit between different random noise levels. This is made possible since our score network is trained without taking random noise’s standard deviations as inputs, unlike [34]. We exponentially reduce the random noise’s standard deviation from the initial value *σ*_1_ to its minimum *σ*_*K*_, instead of taking *K* discrete values:

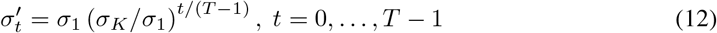

and the update rule for refined 3D coordinates is given by:

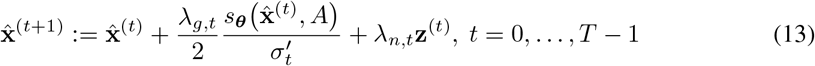

where 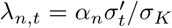 and 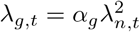. The overall structure optimization process for ALD and CALD is as summarized in Algorithm 1 and 2, respectively. As demonstrated in later experiments, the CALD sampling algorithm leads to more efficient structure optimization than ALD.

#### Algorithm 1 ALD Sampling [34]

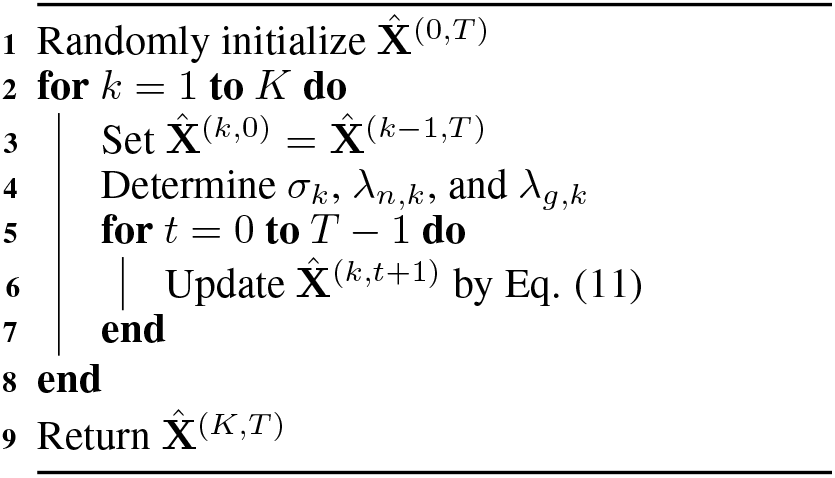

#### Algorithm 2 CALD Sampling (Ours)

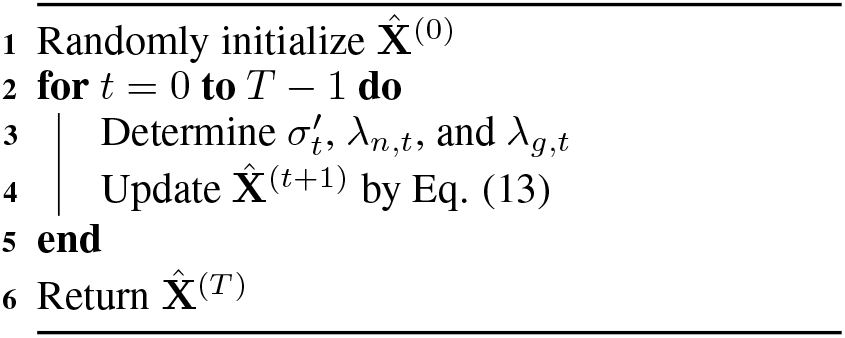

### 4.4 Theoretical Analysis

In this section, we present the theoretical analysis for our SE(3)-Fold’s sampling process. To start with, we show that under mild assumptions, a restrained CALD process is guaranteed to generate samples close to the native protein structure for any amino-acid sequence.

#### Theorem 1.

*Let A be an amino-acid sequence of length L, and* **x**_0_, **x**_1_, … *be a sequence of gradually refined structures, generated by iteratively applying the following update rule:*

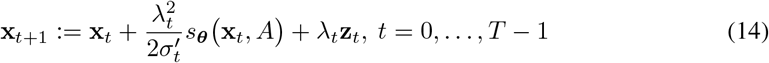

*where* 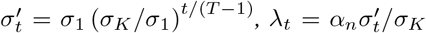, *and* **z**_*t*_ ∼*𝒩* (**0, I**) *is the random noise. With a proper choice of α*_*n*_, *this sequence converges to a probability distribution as T*→ ∞, *whose density function is given by:*

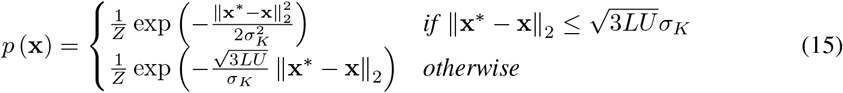

*where Z is the partition constant and* **x**^∗^ *denotes the native protein structure corresponds to the amino-acid sequence A*.

The detailed proof can be found in Appendix A. Theorem 1 indicates that by limiting *α*_*g*_ = 1, the continuously-annealed Langevin dynamics (CALD) sampling process gradually refines 3D coordinates towards a neighborhood of the native protein structure. Furthermore, if the *α*_*g*_ = 1 constraint is discarded, as in the standard CALD sampling process, we show that generated samples will follow a slightly modified probability distribution:

#### Corollary 1.

*If the sequence* **x**_0_, **x**_1_, … *is iteratively generated by:*

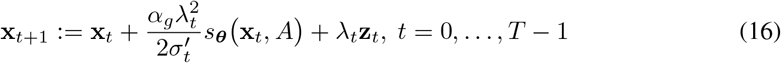

*then this sequence converges to a probability distribution, whose density function is given by:*

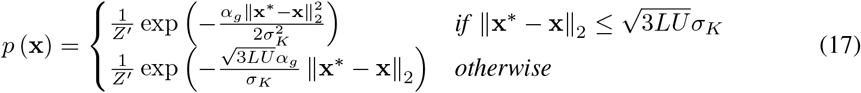

*where Z*′ *is the partition constant, as T* → ∞.

The above probability distribution shares a similar landscape with the original one given in Theorem 1, but the variance can be either increased or decreased, depending on the choice of *α*_*g*_. In other words, this allows us to flexibly adjust the diversity of SE(3)-Fold’s structure optimization results, even though the underlying score network remains the same.

## 5 Experiments

In this section, we firstly report the protein structure optimization accuracy of our proposed SE(3)-Fold method. Afterwards, we demonstrate that our method is computationally efficient and conduct detailed visualization on SE(3)-Fold’s structure optimization process.

### 5.1 Setup

#### Dataset

We randomly select 2065 domains from the CATH database [32] (daily snapshot: 20201021), and split them into training, validation, and test subsets of size 1665/200/200. Particularly, to validate the generalization ability of structure optimization methods, it is guaranteed that each subset has a disjoint set of CATH super-families, *i.e*. no super-family overlap between subsets.

#### Features

For node features, we associate each residue with its amino-acid type’s one-hot encoding and positional encoding to represent its relative position in the amino-acid sequence. For edge features, we employ an inter-residue distance and orientation predictor [42] to infer interactions for all the residue pairs from multiple sequence alignments (MSAs). These predictions are also fed into the baseline structure optimization method for a fair comparison. Please refer to Appendix C for more details on the extraction of node and edge features.

#### Graph Construction

Given a protein, we construct its graph representation with three types of neighbors for each residue: 1) sequence-based neighbors, 2) coordinate-based neighbors, and 3) random neighbors. Sequence-based neighbors connects two residues if their sequence separation is exactly 2^*m*^(*m* = 0, 1, …). Coordinate-based neighbors connects two residues if their C_*α*_-C_*α*_ distance is within top-k minimal ones. Random neighbors introduce randomized connections between residues, which is critical for the sampling process since protein structures are randomly initialized and coordinate-based neighbors at early stages are noisy and less informative.

#### Implementation Details

To verify the effectiveness of multi-head attention, we train two series of score networks for SE(3)-Fold, with either 4 EGCL or MhaEGCL modules. Layer normalization [4] are inserted between EGCL/MhaEGCL modules to normalize intermediate node embeddings. Each model is trained with the Adam optimizer [19] for 50 epochs with batch size 32. The random noise scheme is set as *σ*_1_ = 10.0, *σ*_*K*_ = 0.01, and *K* = 61. We maintain the exponential moving average (EMA) of score network’s model parameters with the momentum parameter *m* set to 0.999, as suggested in [35], and use this EMA-based model for sampling. Hyper-parameters for the sampling process are tuned on the validation subset: *α*_*n*_ = 0.005, *α*_*g*_ = 10, *T* = 16 for ALD, and *T* = 500 for CALD. To alleviate randomness in reported results, we train multiple models with various hyper-parameter combinations, and report the averaged performance of top-10 models (in terms of validation loss) for the subsequent sampling process. The detailed hyper-parameter setting and per-model structure optimization results can be found in Appendix D. The training and sampling of each model is conducted in a single node, with Intel Xeon 8255C CPU and Nvidia Tesla V100 GPU.

### 5.2 Structure Optimization Accuracy

In Table 1, we report averaged and maximal lDDT-Ca scores [22] of several variants of our SE(3)-Fold approach on the test subset. For each model, we repeat the sampling process for 4 times (batch size: 32) to generate 128 decoys per domain. As a comparison, we run trRosetta [48] with the same inter-residue distance and orientation predictions, and generate 300 decoys per domain for evaluation, following its default hyper-parameter setting.

**Table 1:** Comparison on averaged and maximal lDDT-Ca scores of trRosetta and SE(3)-Fold. For trRosetta and SE(3)-Fold-Backbone, lDDT scores for backbone atoms are reported within brackets.

| Method | Format | Layer | Sampling | Avg. IDDT-Ca | Max. IDDT-Ca |
| --- | --- | --- | --- | --- | --- |
| trRosetta [48] | - | - | - | 0.7456 (0.7365) | **0.8525 (0.8465)** |
| EBM-Fold [41] | $C_\alpha$-trace | - | - | 0.7775 | 0.8124 |
| SE(3)-Fold | $C_\alpha$-trace | EGCL | ALD | $0.7545 \pm 0.0089$ | $0.7818 \pm 0.0057$ |
| | | | CALD | $0.7654 \pm 0.0112$ | $0.7965 \pm 0.0054$ |
| | | MhaEGCL | ALD | $0.7796 \pm 0.0065$ | $0.8028 \pm 0.0052$ |
| | | | CALD | $0.7902 \pm 0.0048$ | $0.8134 \pm 0.0037$ |
| | Backbone | MhaEGCL | CALD | **$0.7928 \pm 0.0036$** | $0.8148 \pm 0.0033$ |
| | | | | **($0.7611 \pm 0.0107$)** | ($0.7806 \pm 0.0115$) |

From Table 1, we discover that both multi-head attention-based EGCL (MhaEGCL) module and continuously-annealed Langevin dynamics (CALD) consistently improve the structure optimization accuracy over their counterparts. In general, SE(3)-Fold achieves higher averaged lDDT-Ca scores than trRosetta, but is inferior to it in terms of maximal lDDT-Ca scores. This is mainly due to the notable difference in the decoy quality distribution between trRosetta and SE(3)-Fold. In Figure 1, we visualize the lDDT-Ca distribution for a few test domains, using the best MhaEGCL-based model (in terms of the validation loss) and CALD sampling. It is quite obvious that SE(3)-Fold’s lDDT-Ca distribution is more concentrated than that of trRosetta, indicating that SE(3)-Fold tends to generate more consistent structure predictions. However, the best decoy’s quality of SE(3)-Fold is indeed inferior to trRosetta, which needs further investigation and improvement.

**Figure 1:**
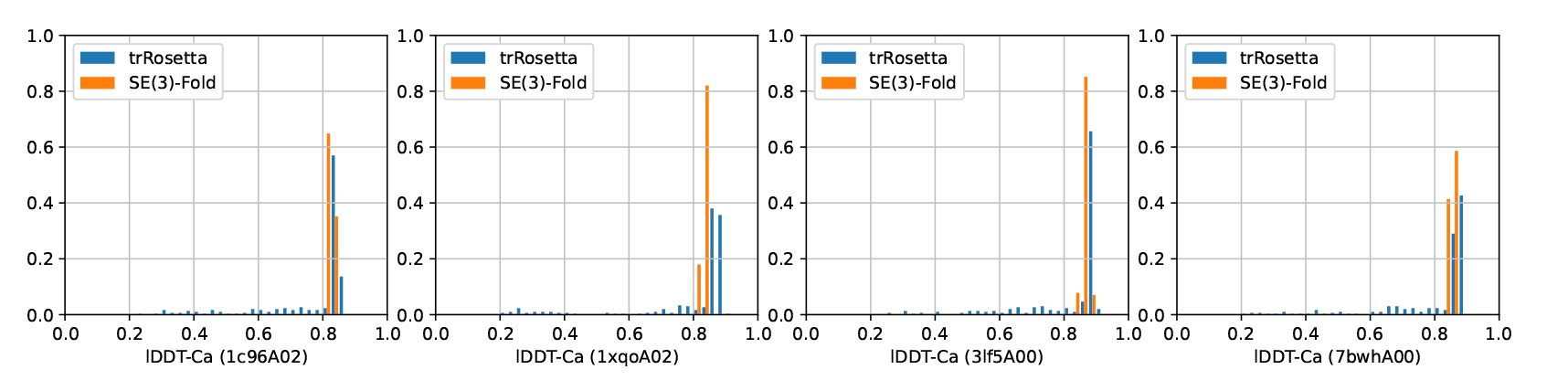
Comparison on lDDT-Ca distributions of trRosetta and SE(3)-Fold (residue format: C_*α*_-trace). lDDT-Ca scores are discretized into 40 bins, and the domain ID is noted in the bracket.

### 5.3 Computational Efficiency

Most of state-of-the-art protein structure optimization algorithms [48, 30, 17] are developed on top of Rosetta software suite [33], which heavily relies on CPU-based computation while general GPU support is not yet presented. In contrast, SE(3)-Fold’s computation overhead is dominated by the score network’s forward pass, which can be efficiently accelerated by modern deep learning toolkits with full GPU support.

In Figure 2, we compare trRosetta and SE(3)-Fold’s structure optimization time for each test domain. For CPU-based optimization, we limit the number of threads to 1 for a fair comparison. For GPU-based implementation, we set the batch size to 32 to improve the GPU utilization, and report the per-decoy optimization time (total time divided by the batch size). Here, we report the time consumption of the network architecture with highest computational overhead among all the candidates.

**Figure 2:**
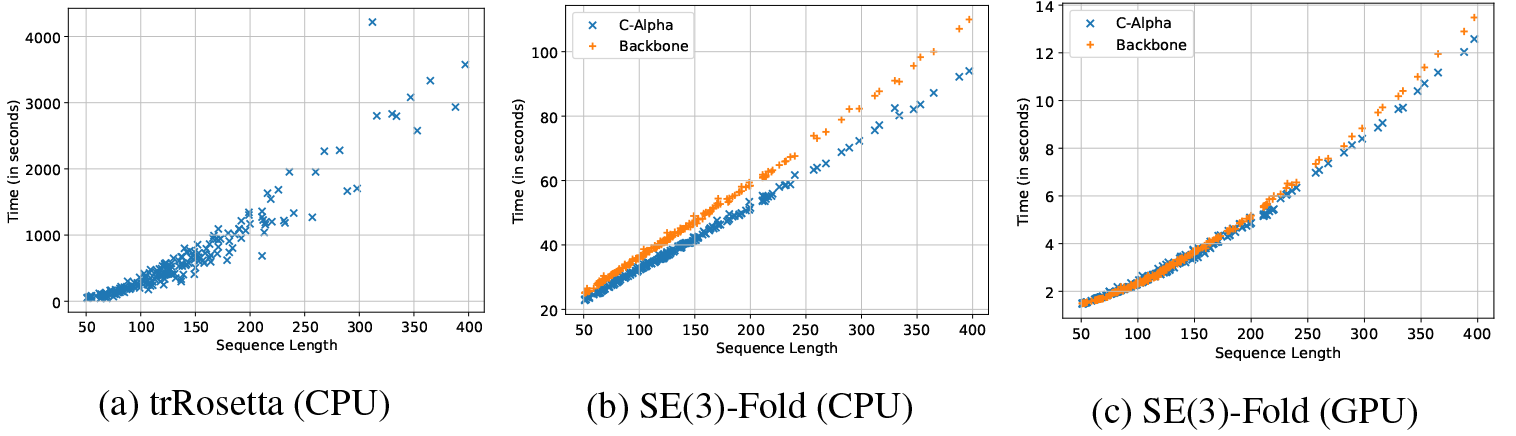
Comparison on the structure optimization time (per decoy) of trRosetta and SE(3)-Fold. For trRosetta, we report the averaged time consumption of three independent runs.

As depicted in Figure 2a, trRosetta’s structure optimization time grows significantly for proteins with longer amino-acid sequences. For 1ojjA00 (length: 397 residues), it takes approximately 1 hour to generate one decoy with trRosetta. On the other hand, SE(3)-Fold’s structure optimization time grows much slower (approximately linear) *w.r.t*. the sequence length. Under the same hardware specification, CPU-based SE(3)-Fold achieves up to one magnitude speed-up over trRosetta, especially for long proteins. With GPU acceleration enabled, it only takes ∼ 13 seconds for SE(3)-Fold to generate one decoy for the longest test domain, which is over 100 times faster than trRosetta. It is also worth mentioning that backbone-based residue format does not leads to much additional time consumption, compared against C_*α*_-atom based representation, especially for GPU-based implementation. Detailed analysis on the computational complexity of SE(3)-Fold can be found in Appendix E.

### 5.4 Visualization on the Sampling Process

Finally, we delve into SE(3)-Fold’s structure optimization process to understand how protein structures are gradually refined from random initialization. In Figure 3, we visualize the lDDT-Ca score *vs*. number of iterations in ALD and CALD sampling process for a few test domains. Both ALD and CALD converge with a similar speed, but CALD avoids zig-zag patterns in ALD, due to discontinuous transition between different random noise levels. Please note that we reduce *T* = 16 to *T* = 8 in ALD to approximately match the total number of iterations with CALD, although *T* = 16 also leads to similar results in ALD.

**Figure 3:**
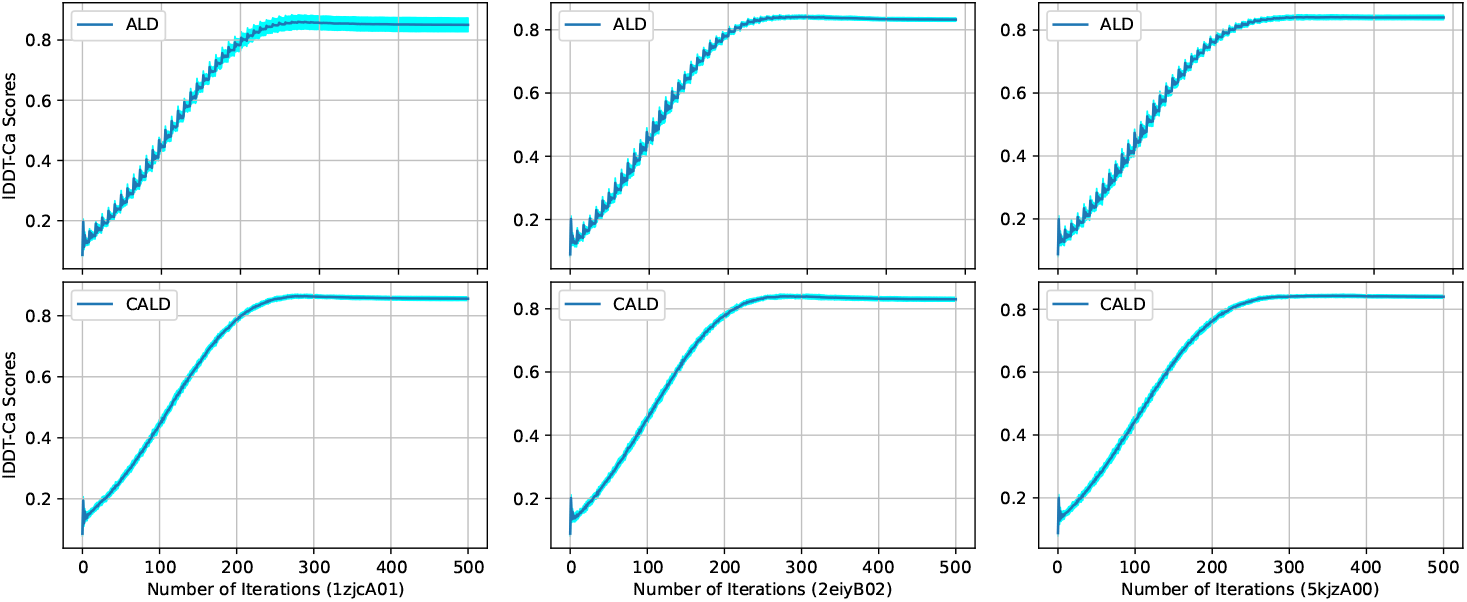
Comparison on lDDT-Ca scores during ALD (top) and CALD (bottom) sampling process. Each column corresponds to the same test domain, with the domain ID noted in the bracket.

## 6 Discussion

In this paper, we propose SE(3)-Fold as a fully-differentiable approach for protein folding. SE(3)-Fold achieves comparable structure optimization accuracy with state-of-the-art Rosetta-based protocols, and is efficient in both training (within one GPU day) and sampling (1-2 orders of magnitude faster than trRosetta). The co-evolution information from homologous sequences are well utilized, which leads to more accurate predictions than other end-to-end approaches, *e.g*. RGN and NEMO.

### Limitations

SE(3)-Fold still relies on a pre-trained network to predict inter-residue distance and orientations from homologous sequences. Such network may be sub-optimal as its optimization does not fully utilize the information from 3D structures. Joint learning of these two modules may further boost the structure optimization accuracy.

## Appendix A Proof for Theorem 1

In Theorem 1, we consider the following update rule:

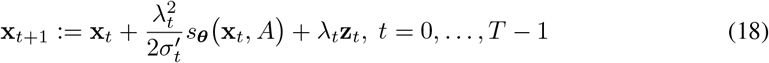

where 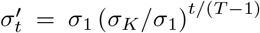 and 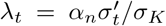. The dimensionality of **x**_*t*_ is denoted as *n* = 3*LU*, which corresponds to an amino-acid sequence of length *L* and *U* atoms per amino-acid residue. We now show that under mild assumptions, the above iterative process converges to a neighborhood of the native protein structure for any specified amino-acid sequence.

To start with, we consider the following oracle score function, which is used to define the probability distribution we wish to converge to:

### Definition 1.

*For an amino-acid sequence A, its oracle score function is defined as:*

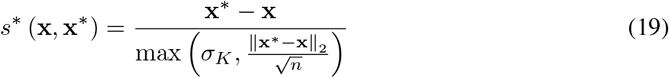

*where* **x**^∗^ *represents the native protein structure, and σ*_*K*_ *is the minimal standard deviation of random noise used during the training process*.

### Lemma 1.

*The oracle score function s*^∗^ () *defines a probability distribution, whose probability density function is given by:*

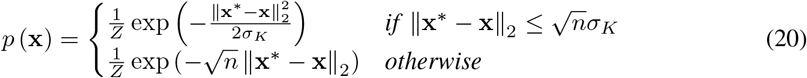

*where Z is the partition constant to guarantee that p* (**x**) *is valid*.

### Proof.

Due to the existence of max operator, we rewrite the oracle score function’s definition as:

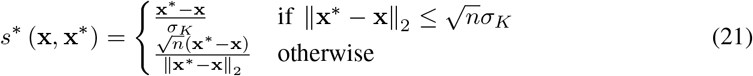

For the first case, we have:

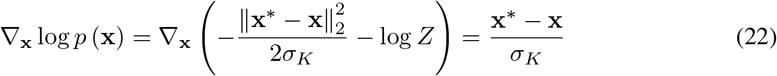

and for the second case, we have:

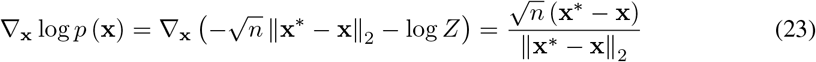

which exactly matches the definition of oracle score function. Next, we show that these exists a partition constant *Z* satisfying that *p* (**x**) is a valid probability distribution. Particularly, we need to prove that the following two integral terms are finite:

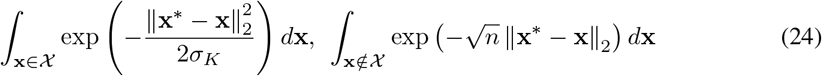

where 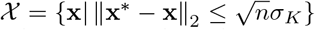. Since the exponential term term only rely on the Euclidean distance between **x** and **x**^∗^, these integral terms can be transformed as:

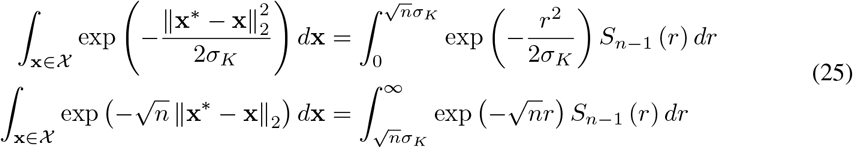

where *r* =‖ **x**^∗^ *−* **x** ‖_2_ and

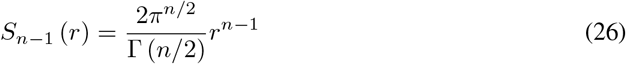

is the surface area of the (*n* − 1)-sphere of radius *r* embedded in the *n*-dimensional space, and Γ (·) is the gamma function. The first integral term is finite since the function within the integral symbol is finite. For the second integral term, we have:

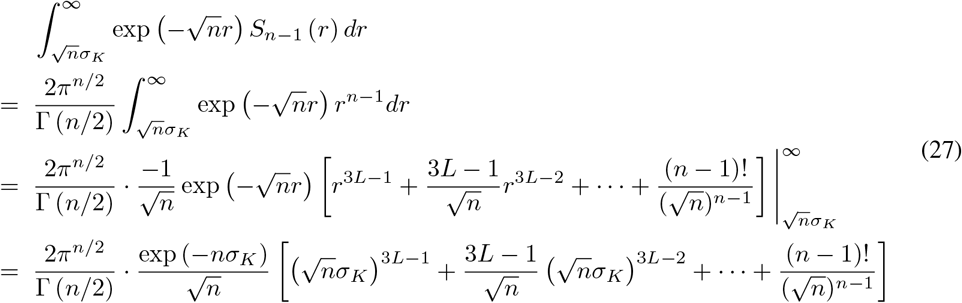

which is also finite. □

### Lemma 2.

*If the oracle score function is re-scaled by σ*_*K*_, *i.e*.:

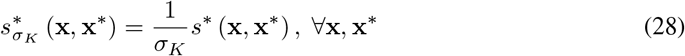

*then it corresponds to a probability distribution, whose probability density function is given by:*

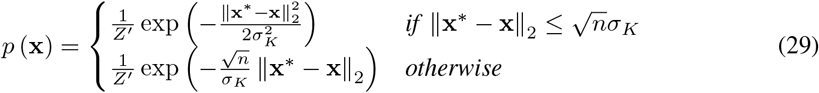

*where Z*′ *is the partition constant*.

It is worthy noting that for both the oracle score function and its re-scaled counterpart, the corresponding probability distribution’s density function has a single global-maximal at **x**^∗^.

During the training phase, we firstly select a random noise level *σ*_*k*_ from *σ*_1_, …, *σ*_*K*_, and then perturb the native protein structure **x**^∗^ with some random noise drawn from 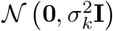. Therefore, the relative difference between native and perturbed structures is approximately given by:

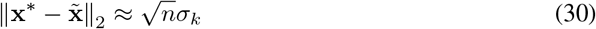

where 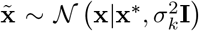. With the above approximation, we can rewrite the original objective function in Eq. (10) as:

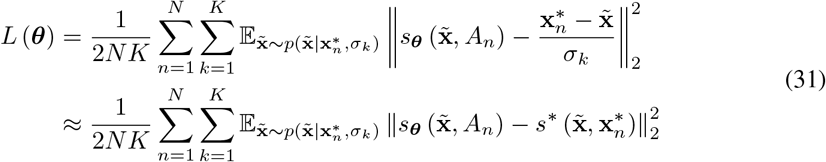

which implies that the score network *s*_***θ***_ (*·*) is essentially approximating the oracle score function during the training process. Additionally, we make following assumptions:

### Assumption 1.

*The score network has bounded variance:*

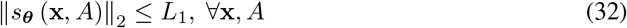

### Assumption 2.

*The score network is Lispchitz continuous:*

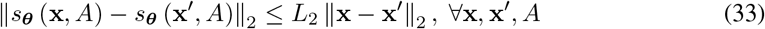

### Assumption 3.

*The score network has bounded estimation error for the oracle score function:*

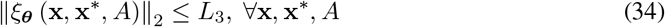

where *ξ* _***θ***_ (x, x^∗^,*A*) = *s*_***θ***_ (x,*A*) − *s*^∗^ (x, x^∗^).

With above analyses and assumptions, we now present the convergence guarantee for our proposed continuously-annealed Langevin dynamics (CALD) sampling process:

### Theorem 1.

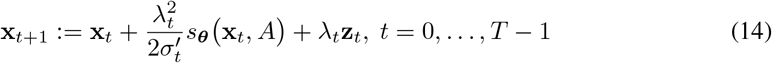

*where* 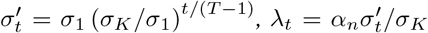, *and* **z**_*t*_ ∼*𝒩* (**0, I**) *is the random noise. With a proper choice of α*_*n*_, *this sequence converges to a probability distribution as T* → ∞, *whose density function is given by:*

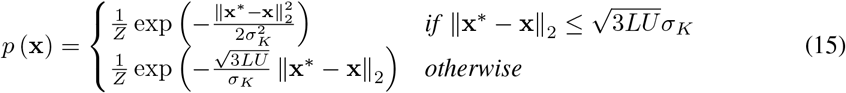

### Proof.

In order to prove the convergence of sequence **x**_0_, **x**_1_, …, we will show that one of its sub-sequences, 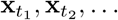, converges to the above probability distribution, therefore the whole sequence also converges.

For any small constant *ϵ* (0 *< ϵ* ≪ 1), with a sufficiently small *α*_*n*_, we can find a sub-sequence *t*_1_ *< t*_2_ *<* … satisfying that:

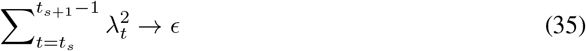

as *s* → ∞.

The overall update process from 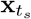 to 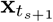 is given by:

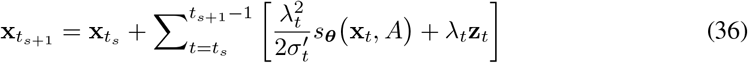

where the update signal essentially consists of two parts: accumulated estimated gradients and random noise. For the latter one, since each iteration’s random noise is independent, we have:

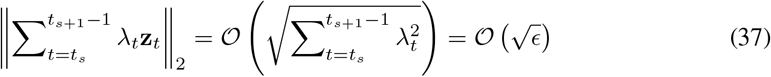

Since the score network is assumed to have bounded variance, for any intermediate iteration *t*′ (*t*_*s*_ *< t*′ ≤ *t*_*s*+1_), the difference between **x**_*t′*_ and 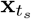 is bounded by:

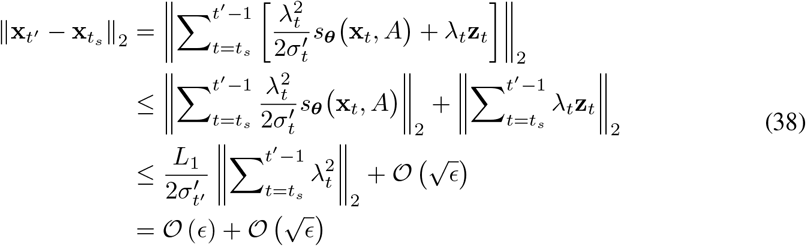

which implies that the difference between the score network’s outputs is also bounded:

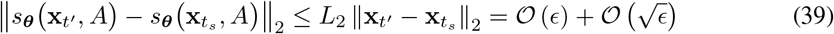

Therefore, for the accumulated score network’s outputs in Eq. (36), we have:

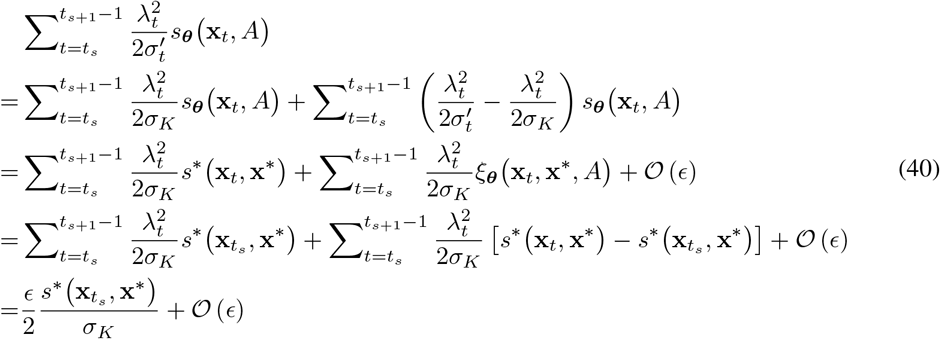

where the second equality holds due to the bounded variance assumption, the third equality holds due to the bounded estimation error assumption, and the last equality holds due to the Lipschitz continuous property of oracle score function, based on its definition.

Hence, the iterative update process from *t*_*s*_ to *t*_*s*+1_ is essentially one gradient step at 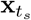 with step size *ϵ* and re-scaled oracle score function, with a deviation term dominated by *𝒪* (*ϵ*) and random noise whose standard deviation equals 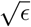. The *𝒪* (*ϵ*) deviation term can be omitted due to the existence of random noise. Therefore, we have:

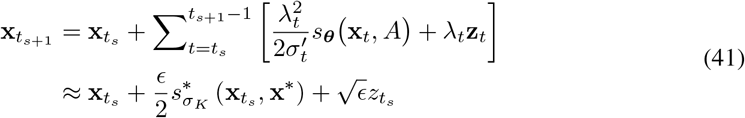

The sub-sequence 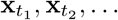 can be viewed as a sequence generated by Langevin dynamics with a constant step size of *ϵ*, which converges to the probability distribution defined by the re-scaled oracle score function, as *ϵ* → 0 and *T* → ∞. According to Lemma 2, the re-scaled oracle score function’s probability distribution is exactly the one as defined in Theorem 1, which concludes the proof. □

## Appendix B Network Architecture

In MhaEGCL, multiple sub-networks are employed to compute edge-wise messages, query and key embeddings, attention coefficients, and weighting coefficients for estimated gradients. In our experiments, we implement these sub-networks with shallow MLP models, as listed in Table 2. The output dimension of each linear layer is given in the bracket, where *D*_*m*_ = *D*_*a*_ = *D*_*o*_ = 16. Three types of non-linear activation functions are used: Swish [25], ELU [6], and LeakyReLU [21].

**Table 2:** Network architectures in the MhaEGCL unit.

| Network | Architecture |
| --- | --- |
| $\phi_m^l$ | Linear ($D_m$) - Swish - Linear ($D_m$) |
| $\phi_q^l$ | Linear ($D_a$) - ELU - Linear ($D_a$) |
| $\phi_k^l$ | Linear ($D_a$) - ELU - Linear ($D_a$) |
| $\phi_e^l$ | Linear (1) - LeakyReLU |
| $\phi_h$ | Linear ($D_o$) - Swish - Linear ($D_o$) |
| $\phi_{s,u}^l$ | Linear ($D_m$) - Swish - Linear (1) |

## Appendix C Node and Edge Features

### Node features

Each node in the graph corresponds to a single amino-acid residue in the protein. We extract two types of node features: one-hot encoding of amino-acid types and positional encoding. For the latter one, we introduce a “maximal sequence length” hyper-parameter *L*_*max*_ (although this also works for sequences longer than *L*_*max*_), and extract features as follows:

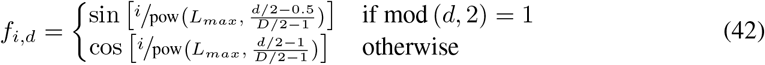

where *i* is the amino-acid residue’s index in the sequence, and *d* = 1, …, *D* denotes the feature index in the *D*-dimensional positional encoding. For all the experiments in this paper, we set *L*_*max*_ = 1000 and *D* = 24.

### Edge features

Each edge in the graph corresponds to a pair of amino-acid residues. As a commonly-used representation for inter-residue interactions, we adopt a MSA-based neural network to predict inter-residue distance and orientations [42], following trRosetta’s definitions:

- *d*_*ij*_: Euclidean distance between 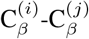 atoms
- *ω*_*ij*_ :dihedral angle defined by 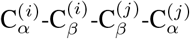 atoms
- *γ*_*ij*_: dihedral angle defined by 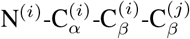 atoms
- *φ*_*ij*_: plane angle defined by 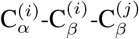 atoms

All of above distance and orientation angles are discretized into histogram bins (37 bins for *d*_*ij*_, 25 bins for *ω*_*ij*_ and *γ*_*ij*_, and 13 bins for *φ*_*ij*_). For each residue pair, we obtain a 100-dimensional feature vector describing their relative distance and orientations, which is then used as edge features.

## Appendix D Detailed Results for Top-ranked Models

For models built with EGCL modules (residue format: C_*α*_-trace), following hyper-parameters are tuned:

- *D*_*m*_ ∈ {16, 32, 64}: number of dimensions for edge-wise messages
- *N*_*s*_ ∈ {4, 8, 12}: number of sequence-based neighbors per residue
- *N*_*c*_ ∈ {4, 8, 12}: number of coordinate-based neighbors per residue
- *N*_*r*_ ∈ {4, 8, 12}: number of random neighbors per residue
- *η ∈* {0.001, 0.003, 0.010}: Adam optimizer’s learning rate

Please note that we limit *N*_*c*_ = *N*_*r*_ to narrow down the hyper-parameter search space. A total of 81 models are trained, from which top-10 models with lowest validation losses are kept for the subsequent sampling process on validation (for hyper-parameter tuning) and test (for final results) subsets. Per-model results are as listed in Table 3.

**Table 3:** Top-10 EGCL-based C_*α*_-trace models’ hyper-parameters, validation losses, and structure optimization results on the test subset (CALD sampling).

| No. | $D_m$ | $N_s$ | $N_c$ | $\eta$ | Loss | IDDT-Ca (Avg.) | IDDT-Ca (Max) |
| --- | --- | --- | --- | --- | --- | --- | --- |
| 1 | 64 | 8 | 4 | 0.001 | 1.3509 | 0.7425 | 0.7858 |
| 2 | 64 | 12 | 4 | 0.001 | 1.3605 | 0.7641 | 0.7941 |
| 3 | 64 | 4 | 8 | 0.001 | 1.3706 | 0.7712 | 0.7986 |
| 4 | 64 | 8 | 8 | 0.001 | 1.3717 | 0.7668 | 0.7932 |
| 5 | 32 | 8 | 4 | 0.003 | 1.3778 | 0.7702 | 0.7995 |
| 6 | 32 | 8 | 8 | 0.003 | 1.3789 | 0.7766 | 0.8021 |
| 7 | 32 | 8 | 8 | 0.001 | 1.3797 | 0.7530 | 0.7943 |
| 8 | 64 | 12 | 8 | 0.001 | 1.3832 | 0.7755 | 0.8033 |
| 9 | 32 | 12 | 8 | 0.001 | 1.3842 | 0.7581 | 0.7928 |
| 10 | 64 | 4 | 12 | 0.001 | 1.3907 | 0.7755 | 0.8008 |

For models built with MhaEGCL modules (residue format: C_*α*_-trace), following hyper-parameters are tuned:

- *N*_*h*_ ∈ {1, 2, 4}: number of attention heads
- *N*_*s*_ ∈ {4, 8, 12}: number of sequence-based neighbors per residue
- *N*_*c*_ ∈ {4, 8, 12}: number of coordinate-based neighbors per residue
- *N*_*r*_ ∈ {4, 8, 12}: number of random neighbors per residue
- *η ∈* {0.001, 0.003, 0.010}: Adam optimizer’s learning rate

Similarly, we limit *N*_*c*_ = *N*_*r*_ to reduce the hyper-parameter search space. We fix *D*_*m*_ = 16 so that the overall model size is similar with EGCL-based models. In Table 4, we report top-10 MhaEGCL-based models’ hyper-parameter settings and structure optimization results.

**Table 4:** Top-10 MhaEGCL-based C_*α*_-trace models’ hyper-parameters, validation losses, and structure optimization results on the test subset (CALD sampling).

| No. | $N_h$ | $N_s$ | $N_c$ | $\eta$ | Loss | IDDT-Ca (Avg.) | IDDT-Ca (Max) |
| --- | --- | --- | --- | --- | --- | --- | --- |
| 1 | 4 | 12 | 12 | 0.003 | 1.2841 | 0.7962 | 0.8183 |
| 2 | 4 | 8 | 12 | 0.003 | 1.2932 | 0.7820 | 0.8069 |
| 3 | 4 | 12 | 8 | 0.003 | 1.2988 | 0.7884 | 0.8113 |
| 4 | 4 | 4 | 12 | 0.003 | 1.2997 | 0.7896 | 0.8127 |
| 5 | 4 | 4 | 12 | 0.010 | 1.3046 | 0.7887 | 0.8120 |
| 6 | 2 | 12 | 8 | 0.003 | 1.3095 | 0.7847 | 0.8102 |
| 7 | 2 | 8 | 12 | 0.003 | 1.3120 | 0.7921 | 0.8135 |
| 8 | 4 | 8 | 8 | 0.003 | 1.3126 | 0.7932 | 0.8152 |
| 9 | 4 | 8 | 12 | 0.001 | 1.3164 | 0.7896 | 0.8147 |
| 10 | 2 | 12 | 12 | 0.003 | 1.3167 | 0.7979 | 0.8192 |

For models built with MhaEGCL modules (residue format: backbone), following hyper-parameters are tuned:

- *N*_*h*_ ∈ {1, 2, 4}: number of attention heads
- *N*_*s*_ ∈ {4, 8, 12}: number of sequence-based neighbors per residue
- *N*_*c*_ ∈ {4, 8, 12}: number of coordinate-based neighbors per residue
- *N*_*r*_ ∈ {4, 8, 12}: number of random neighbors per residue
- *λ*_*b*_ ∈ {0.001, 0.01}: weighting coefficient for estimated gradients over non-C_*α*_ atoms
- *η ∈* {0.001, 0.003, 0.010}: Adam optimizer’s learning rate

where *λ*_*b*_ is newly introduced to control non-C_*α*_ atoms’ contribution to the optimization process. We intentionally put more weights on C_*α*_ atoms since 1) they are more critical in determining the overall structure and 2) the radial distance **r**_*ij*_ for computing edge-wise messages only relies on C_*α*_ atoms. Similar as above, we let *N*_*c*_ = *N*_*r*_ and *D*_*m*_ = 16. In Table 5, we report detailed results for top-10 MhaEGCL-based backbone models.

**Table 5:** Top-10 MhaEGCL-based backbone models’ hyper-parameters, validation losses, and structure optimization results on the test subset (CALD sampling).

| No. | $N_h$ | $N_s$ | $N_c$ | $\lambda_b$ | $\eta$ | Loss | IDDT-Ca (Avg.) | IDDT-Ca (Max) |
| --- | --- | --- | --- | --- | --- | --- | --- | --- |
| 1 | 4 | 12 | 12 | 0.010 | 0.003 | 1.2941 | 0.7988 | 0.8201 |
| 2 | 4 | 12 | 12 | 0.001 | 0.003 | 1.2945 | 0.7980 | 0.8186 |
| 3 | 4 | 12 | 8 | 0.001 | 0.003 | 1.2950 | 0.7899 | 0.8129 |
| 4 | 4 | 4 | 12 | 0.001 | 0.003 | 1.3056 | 0.7929 | 0.8151 |
| 5 | 4 | 8 | 12 | 0.010 | 0.003 | 1.3058 | 0.7944 | 0.8165 |
| 6 | 4 | 4 | 12 | 0.010 | 0.003 | 1.3124 | 0.7925 | 0.8147 |
| 7 | 4 | 8 | 12 | 0.001 | 0.003 | 1.3126 | 0.7930 | 0.8155 |
| 8 | 2 | 12 | 12 | 0.001 | 0.003 | 1.3134 | 0.7894 | 0.8105 |
| 9 | 4 | 12 | 8 | 0.010 | 0.003 | 1.3159 | 0.7913 | 0.8147 |
| 10 | 4 | 12 | 12 | 0.010 | 0.001 | 1.3165 | 0.7874 | 0.8094 |

## Appendix E Analysis on the Computational Complexity

For the sake of convenience, we summarize all the notations to be used for analyzing the computation complexity of MhaEGCL units in Table 6. Each symbol’s typical value is given by either the default setting in our experiments or its largest candidate in the hyper-parameter search space.

**Table 6:**
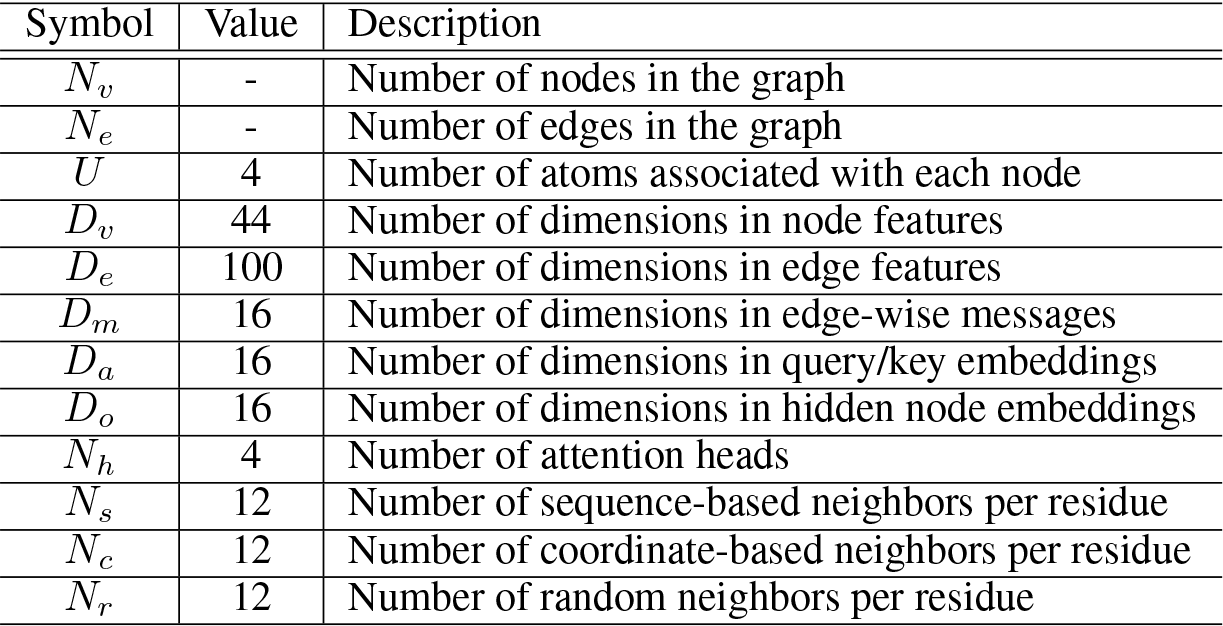
Notations for analyzing the computation complexity of MhaEGCL units.

| Symbol | Value | Description |
| --- | --- | --- |
| $N_v$ | - | Number of nodes in the graph |
| $N_e$ | - | Number of edges in the graph |
| $U$ | 4 | Number of atoms associated with each node |
| $D_v$ | 44 | Number of dimensions in node features |
| $D_e$ | 100 | Number of dimensions in edge features |
| $D_m$ | 16 | Number of dimensions in edge-wise messages |
| $D_a$ | 16 | Number of dimensions in query/key embeddings |
| $D_o$ | 16 | Number of dimensions in hidden node embeddings |
| $N_h$ | 4 | Number of attention heads |
| $N_s$ | 12 | Number of sequence-based neighbors per residue |
| $N_c$ | 12 | Number of coordinate-based neighbors per residue |
| $N_r$ | 12 | Number of random neighbors per residue |

The number of nodes in the graph, *N*_*v*_, equals to the amino-acid sequence length, which is usually within 1000, and the number of edges is given by *N*_*e*_ = (*N*_*s*_ + *N*_*c*_ + *N*_*r*_)*N*_*v*_. For each MhaEGCL unit, we list all the sub-networks’ total computational complexity as below:

- 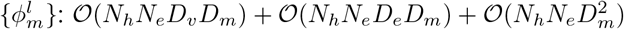
- 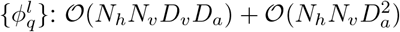
- 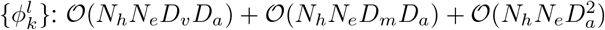
- 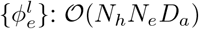
- 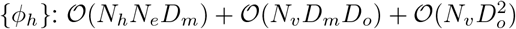
- 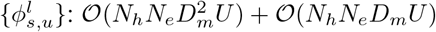

Since the number of dimensions in node/edge features and intermediate embeddings are basically in the same scale, the overall time complexity of one MhaEGCL unit is approximately dominated by 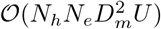. In short, the computational overhead is linear to the number of attention heads, edges in the graph, and atoms associated with each node, and quadratic to the number of dimensions in intermediate embeddings. Since we limit the number of edges per nodes by (*N*_*s*_ + *N*_*c*_ + *N*_*r*_), the overall time complexity is still linear to the number of nodes, *i.e*. the amino-acid sequence length. This is quite favorable, since the time consumption of many Rosetta-based structure optimization methods is quadratic to the sequence length. This is also verified by the comparison on the structure optimization time, as illustrated in Figure 2.

## Appendix F Additional Visualization Results

Here, we take a few test domains as examples, and visualize 2D distance matrices and 3D structures during SE(3)-Fold’s sampling process. For subsequent experiments, we use MhaEGCL-based C_*α*_-trace models with CALD sampling. Since SE(3)-Fold only generates C_*α*_-trace models, we derive corresponding backbone models with PDB_Tool^3^, and then use SCWRL4 [20] to recover side-chain structures.

In Figure 4, 5, and 6, we present visualization on the sampling process for “1zjcA01”, “2eiyB02”, and “5kjzA00”, respectively. We observe that although the initial structure is completely different from the native one (second column), major patterns quickly emerge in the 2D distance matrix within 50-100 iterations. The distance between adjacent residues’ C_*α*_ atoms is not yet correctly preserved at this stage, as indicated by the vast majority of dashed lines in full-atom models’ visualization. After 200 iterations, the overall topology becomes clear and some of secondary structures (*e.g. α*-helices) are already partially visible in C_*α*_-trace and full-atom models. Finally, when 500 iterations are accomplished, almost all the secondary structures are correctly recovered, and the overall topology is also highly similar with the native structure.

**Figure 4:**
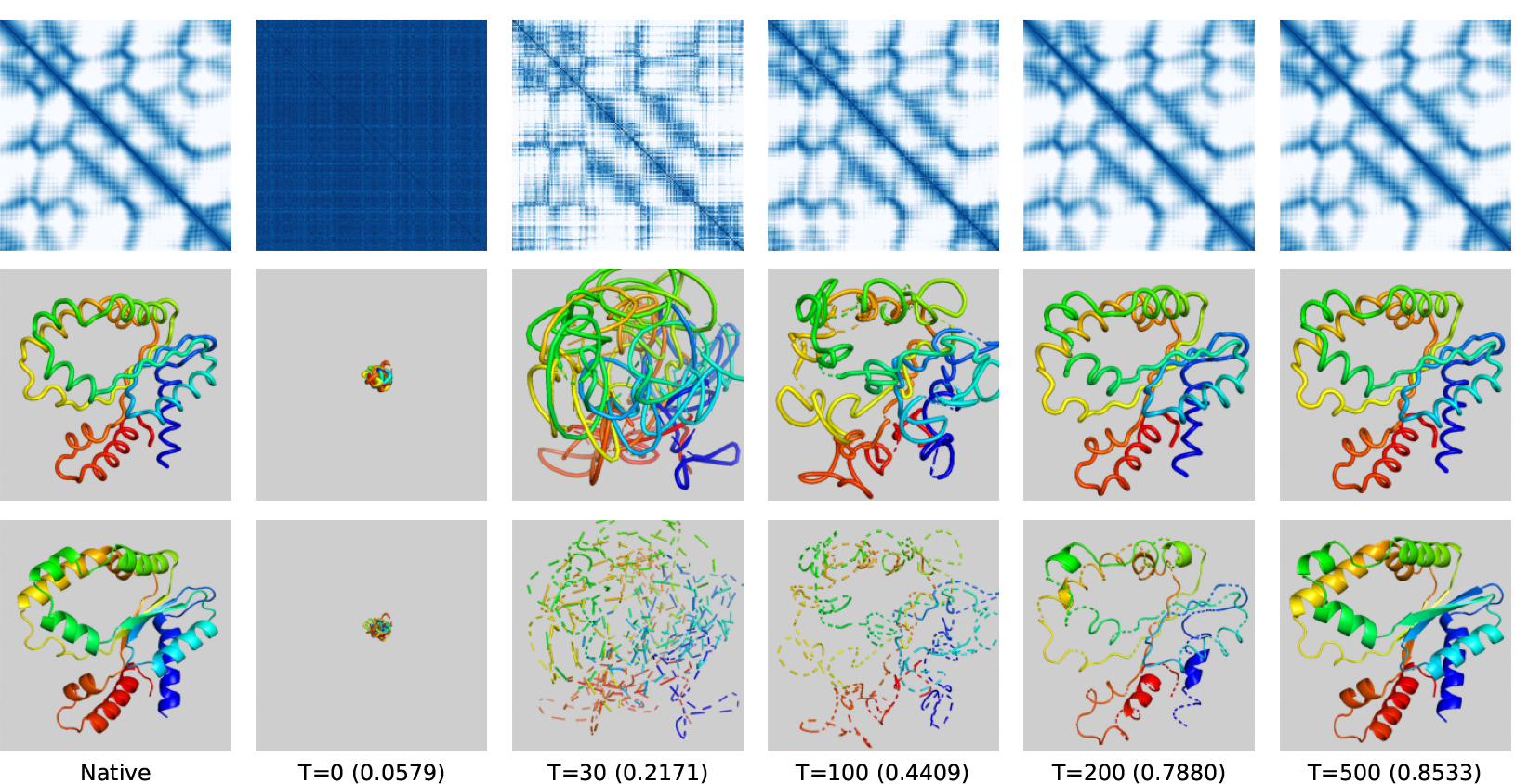
Visualization on SE(3)-Fold’s structure optimization process (CATH domain ID: 1zjcA01). Top: C_*α*_-C_*α*_ distance matrix (darker color corresponds to shorter distance). Middle: C_*α*_-trace models. Bottom: full-atom models recovered with PDB_Tool and SCWRL4.

**Figure 5:**
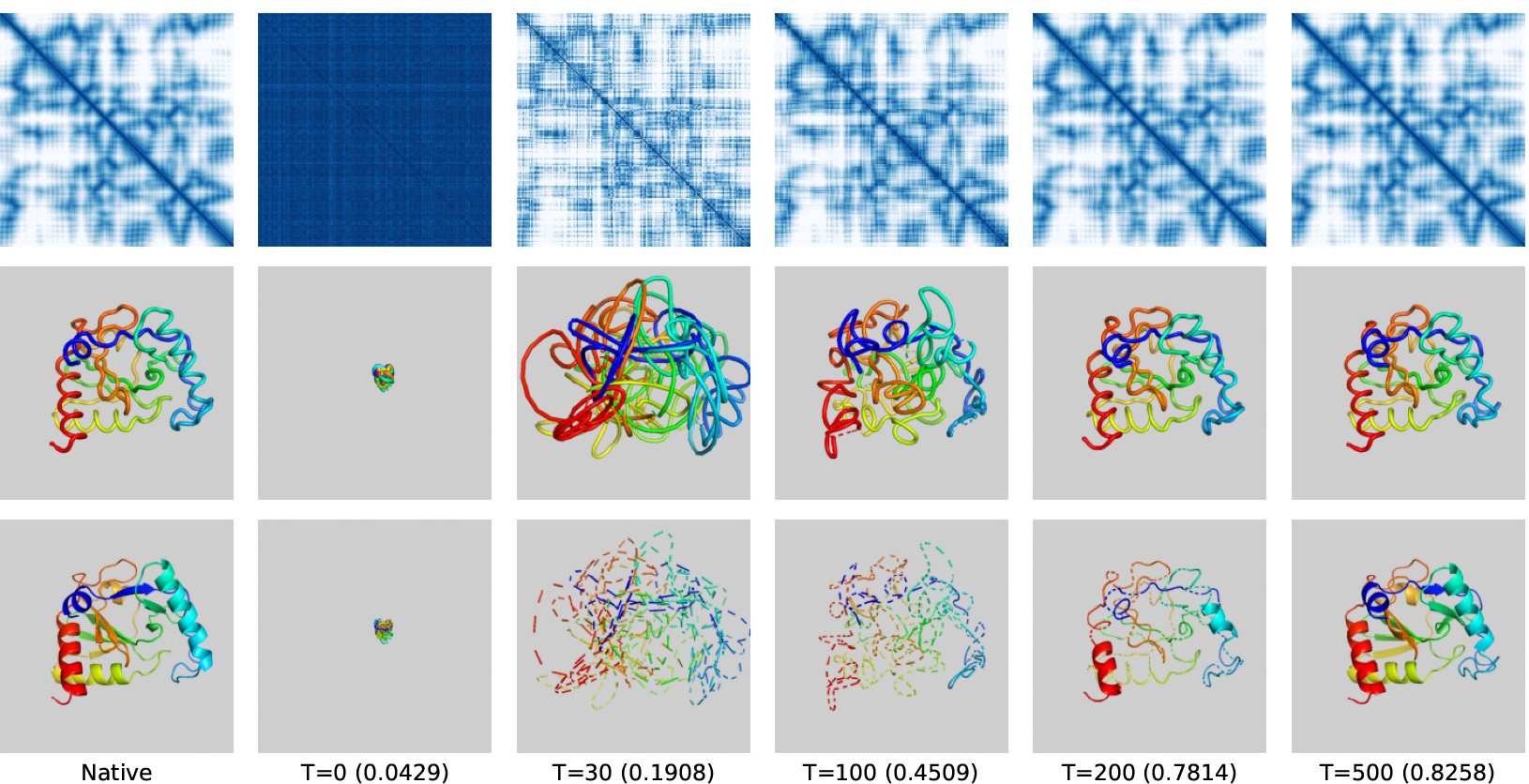
Visualization on SE(3)-Fold’s structure optimization process (CATH domain ID: 2eiyB02). Top: C_*α*_-C_*α*_ distance matrix (darker color corresponds to shorter distance). Middle: C_*α*_-trace models. Bottom: full-atom models recovered with PDB_Tool and SCWRL4.

**Figure 6:**
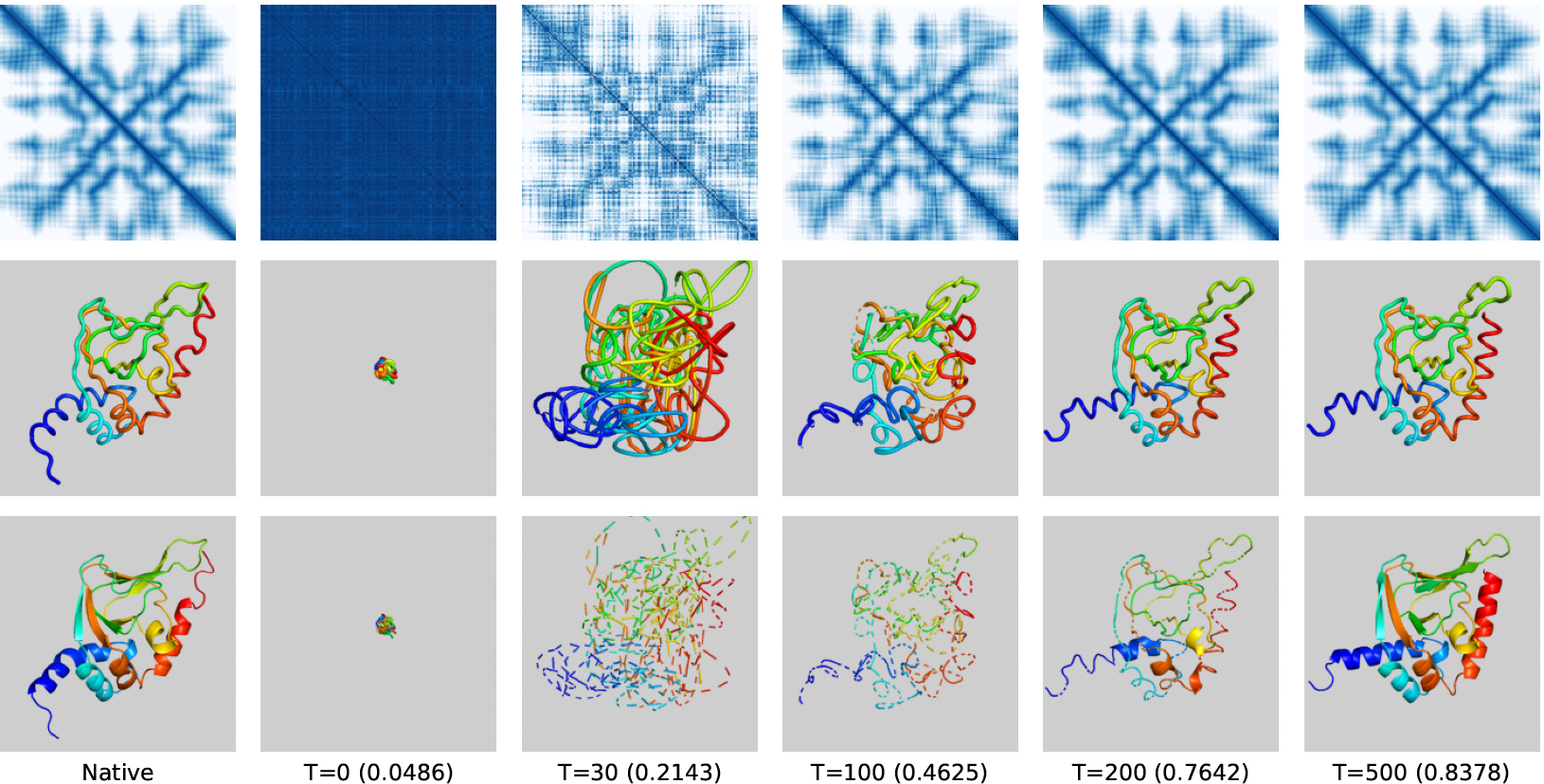
Visualization on SE(3)-Fold’s structure optimization process (CATH domain ID: 5kjzA00). Top: C_*α*_-C_*α*_ distance matrix (darker color corresponds to shorter distance). Middle: C_*α*_-trace models. Bottom: full-atom models recovered with PDB_Tool and SCWRL4.

## Footnotes

1 The CASP14 (Critical Assessment of Structure Prediction) competition, hold on 2020, was recognized as one of the best benchmarks for determining state-of-the-art methods for protein structure prediction.

2 To formulate full-atom structures, *U* = 15 is sufficient to represent all the non-hydrogen atoms in each amino-acid. Zero-padding is required for amino-acids with fewer non-hydrogen atoms.

3 https://github.com/realbigws/PDB_Tool

## Notes

### Competing Interest Statement

The authors have declared no competing interest.

